# Efficient tagging and purification of endogenous proteins for structural studies by single particle cryo-EM

**DOI:** 10.1101/2021.06.25.449985

**Authors:** Jianhua Zhao, Suraj Makhija, Bo Huang, Yifan Cheng

**Author notes:** These authors contributed equally (Jianhua Zhao, Suraj Makhija). Cell and Molecular Biology Program, Sanford Burnham Prebys Medical Discovery Institute, San Diego, CA 92037. Scribe Therapeutics, Alameda, CA 94501.

## Abstract

A major bottleneck in structural biology is producing biologically relevant samples at sufficient quantities. This is particularly true for large protein assemblies where conventional techniques of gene overexpression require substantial optimization, hampering structural studies and drug development efforts. Here we describe a method combining CRISPR/Cas gene editing and fluorescence cell sorting to rapidly tag and purify endogenous human proteins from cell lines, enabling structural analysis of native proteins that are properly folded and assembled. We applied this approach to study the human proteasome from HEK cells and rapidly determined structures of major proteasomal complexes. Structures of the PA28-20S complex reveal the native subunit stoichiometry of PA28 and a distinct functional state of the complex. The efficient strategy for tagging and extracting endogenous proteins described here will enable the structural study of many challenging targets and provide more biologically relevant samples for research and therapeutic development.

## INTRODUCTION

Recent technological breakthroughs in single particle cryogenic electron microscopy (cryo-EM) have greatly accelerated the pace of high-resolution structure determination of challenging biologic macromolecular complexes. However, sample production, namely making properly folded and functional proteins in sufficient quantities, remains one of the biggest limitations in structural biology. Conventional methods of protein production involve over-expression of a target gene from a plasmid within heterologous systems, including bacterial (E. coli), insect (Sf9), and mammalian (HEK) cells. Over-expression approaches frequently encounter issues with proteins that are misfolded, non-functional, or degraded by the host cells. Substantial optimization is often necessary to identify the right conditions to produce suitable proteins for study, which is time-consuming and unsuccessful in many cases. The problem is compounded further by multi-protein complexes where multiple subunits need to be assembled in the right order and at the correct stoichiometries, making the co-expression of multiple subunits and purification of large protein assemblies a major bottleneck to structural studies.

Studies of challenging proteins and large protein complexes have sometimes been possible by extracting samples from natural sources. While this approach have proven successful in special cases, examples of native proteins purified in this way have been largely limited to highly abundant and large protein assemblies that can be isolated by sucrose gradient ultracentrifugation ^1^, or where a known binding partner can be used as bait ^2,3^. A general and efficient method to isolate endogenous proteins from native sources with high specificity and at sufficient quantities for structural studies is currently not available.

Recent technological advances in clustered regularly interspaced palindromic repeats (CRISPR) / Cas-mediated genome engineering have inspired a new way to target endogenous proteins from native sources by genetically incorporating an affinity tag onto a protein of interest for purification and structural analysis ^4,5^. However, the time-consuming process of generating and selecting gene-edited cells have become a barrier to the widespread adoption of this approach. Here we describe a general and highly efficient method to tag endogenous proteins in mammalian cells for purification and structural analysis. Leveraging the unique properties of split fluorescent proteins ^6^, we utilize a small peptide tag that enables both cell selection and affinity purification while minimizing potential structural perturbations.

We demonstrate this approach on the study of endogenous human proteasomal complexes. The proteasome plays an essential role in protein degradation and is critical to maintaining cellular protein homeostasis ^7^. Changes to proteostasis in cells and decreases in protein degradation due to a decline in proteasomal activity have been linked to aging and neurodegeneration ^8^. While the proteasome has been a topic of extensive studies primarily centering on the bacterial homologues, the study of the mammalian proteasome remains challenging due to its structural complexity. Previous studies have utilized complex strategies to capture distinct human proteasomal complexes ^9–12^, but a general strategy to analyze different proteasomal complexes from the same system and at the same time is not available. Here, we demonstrate the ability to visualize and study multiple proteasomal complexes simultaneously by single particle cryo-EM. Additionally, we zoom in on a particular proteasome complex to reveal novel aspects of proteasome function, thus demonstrating the power of the approach described in this study.

## RESULTS

### Rapid tagging and purification of endogenous proteins

A general strategy for tagging and extracting endogenous proteins for structural studies requires three considerations: (1) the tag needs to have a highly specific and high affinity binder for purification, (2) identification and isolation of cells with the correct tag insertion need to be rapid, and most importantly (3) the overall tag must be small to minimize potential structural perturbations and to enable the use of the highly resource-efficient and scalable CRISPR/Cas9 knock-in pipeline 6,13, which has recently been scaled up to tag ~1300 genes in human cells 14. For this purpose, we engineered a tandem FP11-StrepII tag (29 a.a. including the linker, which fits well into the size limit of synthetic single-strand oligo donor DNAs). In this tag, FP11 provides the capability for rapid selection by fluorescence assisted cell sorting (FACS), and StrepII is a small and highly specific marker for purification (Figure 1). FP11’s, including GFP[11] 13, mNeonGreen2[11] 15 and sfCherry2[11] 16, are derived by splitting out the last β strand of fluorescent proteins. Once the tagged protein is expressed, they can spontaneously complement with the corresponding FP1-10 fragment to form a functional fluorescent protein, enabling subsequent sorting of cells containing correct knock-in’s by FACS. Moreover, our method utilizes a FP1-10 fragment that is only transiently expressed, resulting in purified proteins from expanded cells that do not contain a bulky, full-sized fluorescent protein that may interfere with structure or function. The entire process, from design of gRNA and templates to purified protein, can be completed in 3-4 weeks, with multiple experiments easily performed in parallel.

**Figure 1.**
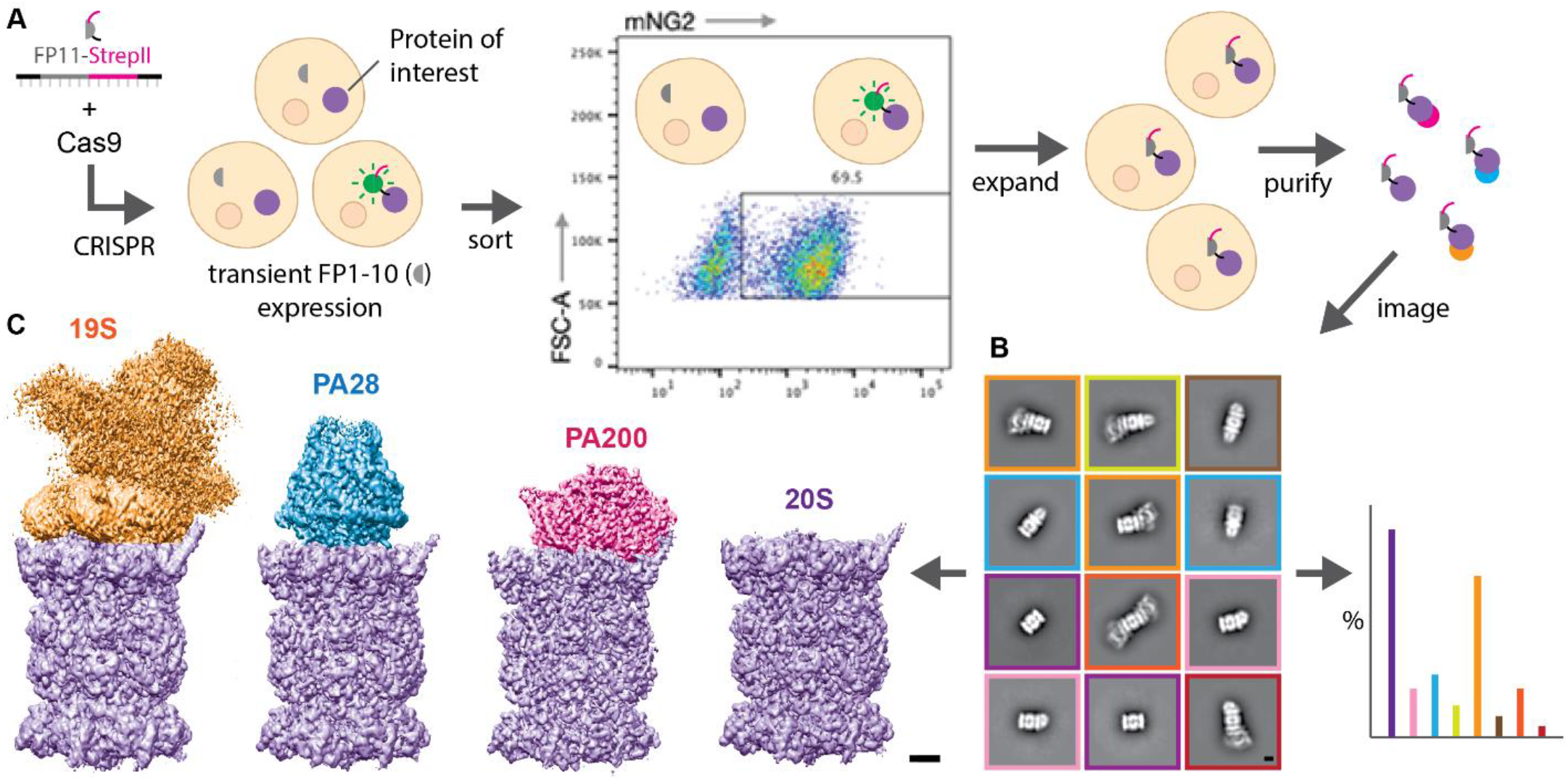
Rapid tagging and purification of endogenous proteins for structural studies. (**A**) Single-stranded DNA encoding FP11-StrepII is delivered into the nucleus of HEK cells along with Cas9 protein complexed with gene-specific guide RNA. Subsequent DNA cleavage and repair results in a mixed population of cells with and without FP11-StrepII successfully incorporated. Transient expression of FP1-10 in the cells enables conjugation between FP1-10 and FP11 to form fluorescent molecules, allowing rapid isolation of cells containing the tag by fluorescence assisted cell sorting (FACS). The isolated cell population can be expanded without the need for single cell clonal selection to allow rapid scaling up of cultures for structural studies. Additionally, the cells are expanded in the absence of FP1-10, resulting in a short 29 amino acid tag on the target protein. The protein can be subsequently purified using the robust and highly specific StrepII affinity marker. (**B**) Tagging and purification of the endogenous human proteasome enables the isolation of a diverse population of different proteasomal complexes, which can be visualized by negative stain EM and 2D classification. Scale bar, 50 Å. (**C**) The purified samples can be analyzed by single particle cryo-EM to determine high-resolution structures of the different complexes. Scale bar, 20 Å.

To demonstrate this approach, we used the human proteasome as a model system. The mammalian proteasome core particle, 20S, is a large barrel-shaped protein complex comprised of four stacked heptameric rings, which include two inner β-rings flanked by two outer α-rings that are each formed by 7 different α or β subunits ^17^. The proteolytic sites are located in the N termini of β-subunits facing interior of the proteolytic chamber. In each α-ring, the N termini of the α subunits form a gate preventing unregulated protein degradation ^18^. There are three known types of endogenous proteasomal activators that bind to the 20S α-ring to open its gate. With two α-rings in 20S, activators can bind in mixed configurations to produce a diverse population with different activators bound. One activator is the ATP-dependent 19S complex that recognizes and unfolds ubiquitinated substrates for degradation in an ATP dependent manner. The function of the other two activators, PA28 and PA200, are ATP independent. PA28 is upregulated by interferon-γ during antigen presentation ^19,20^ and oxidative stress to degrade unfolded proteins ^21^. On the other hand, PA200 is found in the nucleus and is believed to be important for the degradation of histone proteins in DNA repair ^22^.

To facilitate the acquisition of sufficient material for structural studies, Expi293 cells were used to enable high-density growth in suspension. Because the C termini of the β subunits are exposed outside of the 20S barrel, we incorporated a mNG2[11]-StrepII tag onto the C terminus of the 20S β1 (Psmb6), β2 (Psmb7), and β4 (Psmb2) subunits to create three different cell lines. FACS sorting of the cells revealed positive knock-in efficiencies of 5%, 11%, and 24% for β1, β2, and β4, respectively (Supplementary Figure 1). While FACS can separate cells with and without successful knock-ins, the different efficiencies highlight the importance of subunit choice and targeting multiple subunits in parallel to maximize success when tagging protein complexes. Focusing on the β4/Psmb2 cell line, we purified the human proteasome in complex with different activators and visualized these assemblies by negative stain EM (Figure 1B
, Supplementary Figure 2). In the absence of additional ATP or its analogs, affinity pull-down of 20S resulted in mostly 20S alone or capped on one end with PA200, allowing us to determine cryo-EM structures of these complexes to ~2.8 and ~3.2 Å resolution, respectively (Figure 1C, Supplementary Figure 3, Supplementary Table 1). The lack of nucleotide during purification likely led to the disassociation of the 19S regulatory cap from 20S. Indeed, purification in the presence of the ATP analog AMP-PNP allowed us to capture and determine a cryo-EM map of the 19S-20S complex, which appears to be an average of different conformations (Figure 1C). The conformational states of the human 19S-20S proteasome have been well characterized recently ^10,11^ and therefore we did not perform detailed analysis of this proteasome assembly.

### Mapping protein interactions at high resolution

The purification of endogenous proteins captures their native interaction partners and allows visualization of the protein complexes that exist in the cell. Computational classification of the cryo-EM images can further help to shed light on the relative abundance of these complexes, enabling the construction of semi-quantitative high-resolution protein interaction maps (Figure 2). The purification of the 20S proteasome with a tag on β4/Psmb2 enabled the isolation of 20S in complex with its expected interaction partners: 19S, PA200, and PA28 (Figure 1). While double-capped proteasomes were visualized in the negative stain analysis of this sample (Supplementary Figure 2), double-capped species were not observed as a significant population in the cryo-EM images (Figure 2A). Some of the protein complexes likely dissociated during cryo-EM sample preparation, suggesting that negative stain EM is potentially more reliable than cryo-EM for quantifying the distribution of different protein assemblies. Despite this potential limitation, the observed proportion of 19S-20S complexes to PA200-20S and PA28-20S complexes is in line with the expected proportions of these complexes in the cell 8,23. Future work would be necessary to enable more robust quantifications of the absolute abundance of protein complexes in a sample, perhaps by cross-linking the proteins to preserve their integrity prior to sample preparation and imaging.

**Figure 2.**
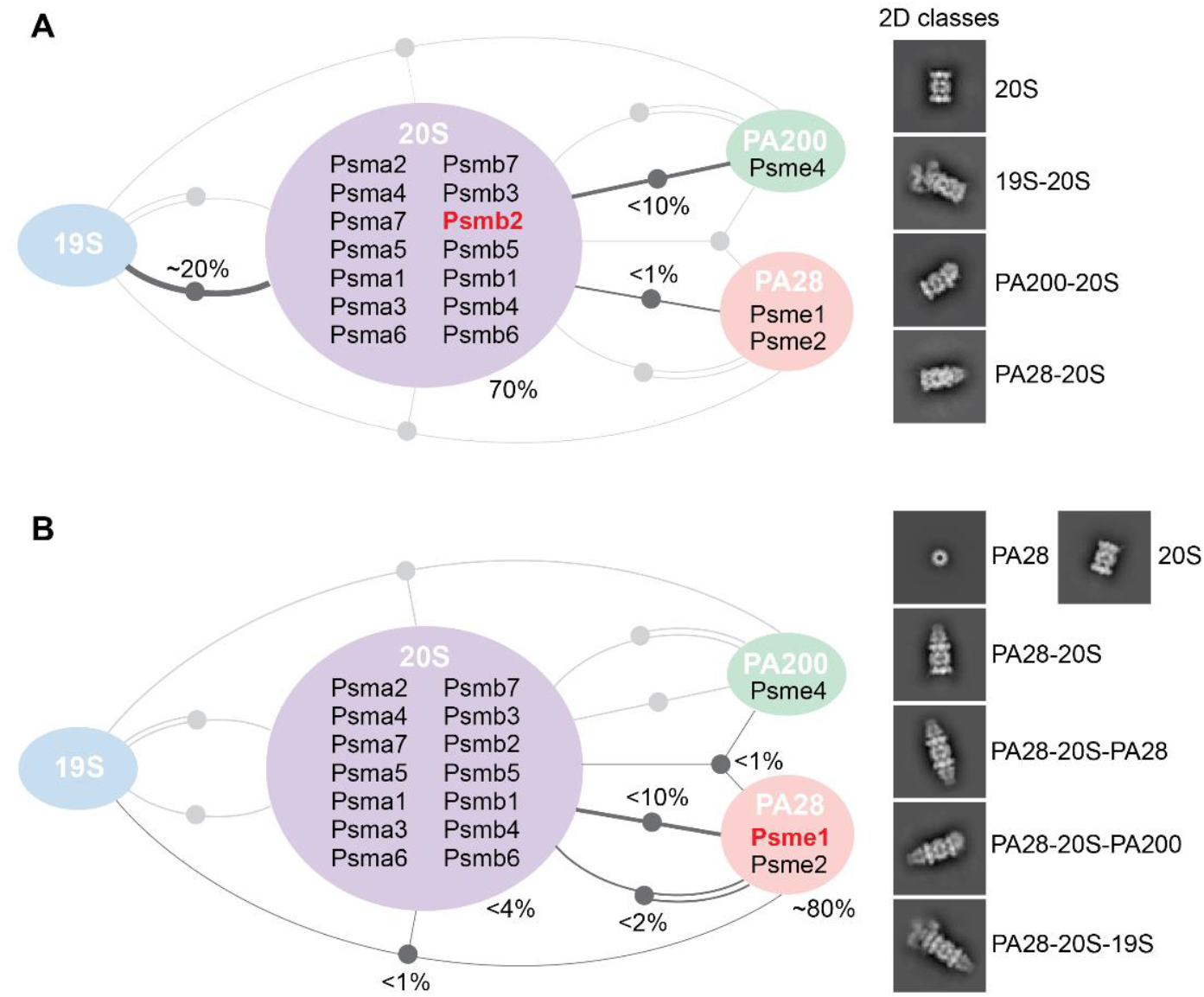
Semi-quantitative protein interaction maps of Psmb2 and Psme1. (**A**) Tagging and affinity pull-down of 20S β4 subunit (Psmb2) co-purifies with activators 19S, PA28, and PA200. Subsequent cryo-EM imaging and computational classification reveals the relative proportion of different complexes. Lines connecting different components indicate an observed interaction (as shown by the 2D images on the right); double lines indicate two copies of a particular activator. Percentages indicate the relative proportion of particle images in a particular class (designated by colored region or node on lines connecting different components). (**B**) Tagging and affinity pull-down of PA28 α subunit (Psme1) enriches for proteasomal complexes containing PA28.

Computational sorting and classification of cryo-EM images allow structure determination of multiple protein assemblies from a heterogenous sample. However, practical restrictions on dataset size impose certain limitations on high-resolution structure determination of complexes with low abundance. Pull-down of 20S by StrepII-tagged β4 (Psmb2) resulted in a mix of different complexes that allowed structure determination of proteasome assemblies making up at least ~10% of the total particle images (Figure 2A). On the other hand, structural analysis of PA28-20S, which comprised less than 1% of the particle images, was challenging. To study this relatively rare class, we engineered an sfCherry2[11]-ALFAtag onto the N-terminus of the PA28 α subunit using the method described above, except here we used a split sfCherry system ^16^ rather than split mNG2. One step affinity purification using an NbALFA nanobody resin ^24^ yielded proteasome assemblies capped with PA28 (Figure 2B), including double-capped proteasomes. Cryo-EM analysis of this sample allowed us to determine structures of single-capped PA28-20S and double-capped PA28-20S-PA28 to ~2.8 and ~3.3 Å resolution, respectively (Figure 3, Supplementary Figure 3, Supplementary Table 1). Thus, we demonstrate the ability to target specific complexes, including rare subpopulations, by tagging select proteins. Additionally, the tagging of PA28 with sfCherry2[11]-ALFAtag was performed in the cell line that already contained mNG2[11]-StrepII-tagged 20S, demonstrating the ability to multiplex orthogonal tags onto different subunits of a larger protein assembly to increase target specificity.

**Figure 3.**
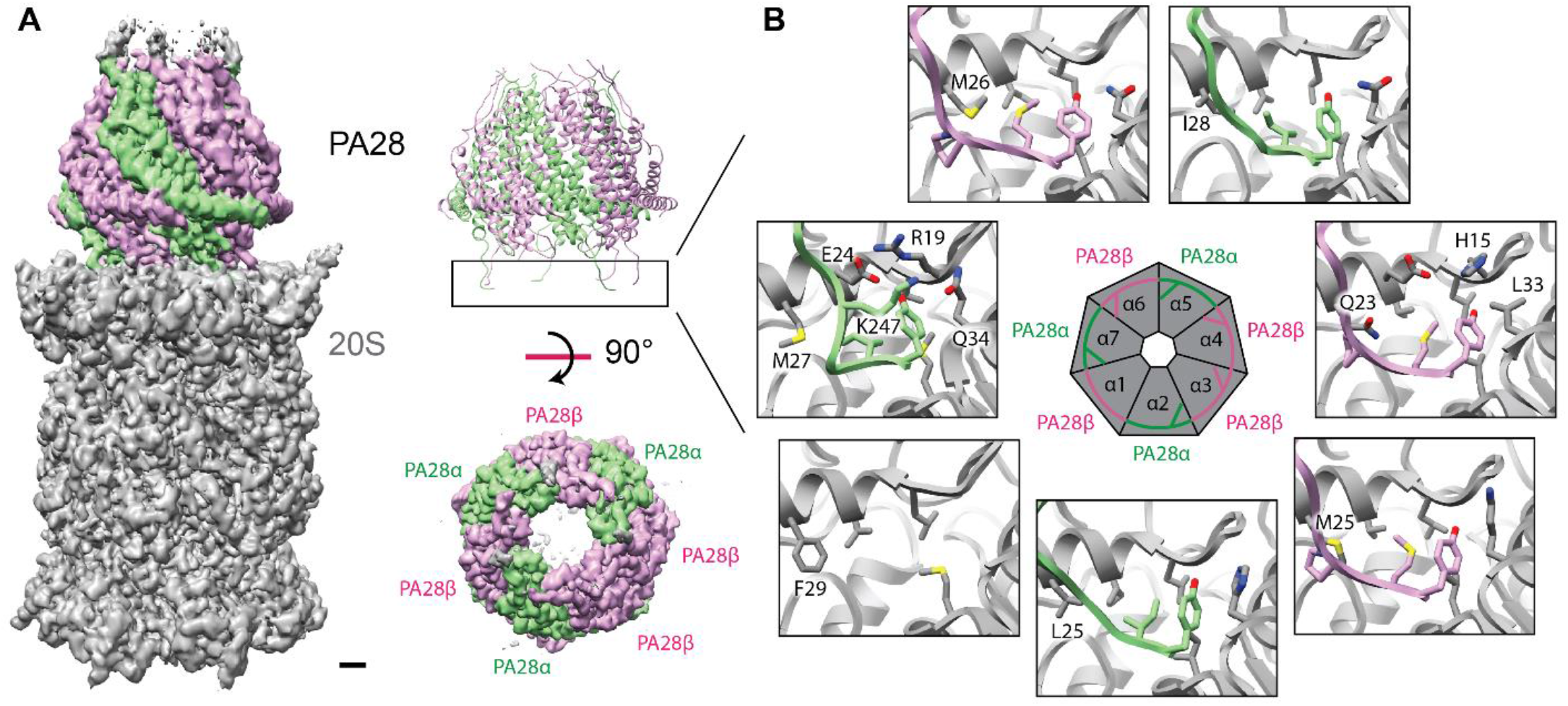
Structure and stoichiometry of the endogenous human PA28-20S proteasomal complex. (**A**) Cryo-EM map of PA28-20S shows PA28 associating tightly with 20S, resulting in well-defined density for the entire complex. The high-quality density map allows clear assignment of 3 α and 4 β subunits in PA28 (Supplementary Figure 4). Scale bar, 10 Å. (**B**) 6 out of 7 binding sites on the α-ring of 20S is occupied by the C-terminal tails of PA28 subunits, providing a strong anchor for PA28 on 20S. The unoccupied binding site located between 20S α1 and α2 contains phenylalanine F29, which is unique at that position and may be hindering the binding of the C-terminal tail of PA28.

### Stoichiometry of the endogenous PA28

Conventional methods of protein overexpression in non-native conditions can result in protein assemblies that do not accurately represent their physiological stoichiometries, which has historically been a challenging problem in structural biology. One advantage of targeting endogenous proteins is it provides macromolecular assemblies in their native configurations, enabling studies that are more biologically relevant. One example that highlights the benefit of this approach is the proteasomal activator PA28, which is comprised of alternating α and β subunits arranged into a heptameric ring with a pseudo-7-fold symmetry. Previous crystallographic structural analysis of the mouse PA28αβ hetero-heptamer found a stoichiometry of 4 α and 3 β subunits ^25^. The protein complex for this previous study was produced by heterologous co-expression of the two subunits, which form a 4α/3β assembly reportedly due to a more thermodynamically stable α-α interface compared to the β-β interface. A more recent cryo-EM study of heterologously expressed human PA28αβ mixed *in vitro* with bovine 20S also made a similar conclusion ^9^. Interestingly, our study of the endogenous human PA28-20S proteasome revealed a different stoichiometry of PA28. The high-quality density map of the PA28-20S complex reported here provides unambiguous assignment of 3 α and 4 β subunits (Figure 3A, Supplementary Figure 4), different from the 4α/3β assembly observed previously.

What might be the functional importance of PA28 stoichiometry? Our structure of the PA28-20S complex shows six out of seven C termini of PA28 engaged in stable interactions with the binding pockets of the 20S alpha-ring (Figure 3B). There is one low-occupancy binding site situated between the 20S α1 and α2 subunits where a phenylalanine, α1 F29, may be sterically hindering the binding of the C-terminal tail of the PA28 β subunit (Figure 3B). The other six unique binding pockets are well engaged with the corresponding C termini of the PA28 3α/4β assembly. On the other hand, the PA28 4α/3β assembly derived from heterologous co-expression showed only partial association with native 20S when assembled *in vitro* ^9^. These results suggest that PA28 in a 4α/3β arrangement has reduced binding affinity to 20S compared to a 3α/4β configuration. The PA28(3α/4β)-20S complex is likely the stable species under native conditions that facilitates the degradation of oxidized and unfolded proteins in the cell.

### Protein loading into the PA28-20S proteasome

Studying endogenous proteins provides a glimpse into their function under native settings and can offer new insights into their mechanisms of action. While it has been well established that the 20S proteasome is a barrel-shaped protein complex having three inner chambers, the function of this modular architecture is not well understood. The catalytic chamber in the middle of the protein complex contains the proteolytic β subunits that are responsible for cleaving polypeptides, but the function of the two adjacent antechambers is not clear. Interestingly, our cryo-EM map of the PA28-20S assembly shows additional density in the antechamber of 20S that is adjacent to PA28 (Figure 4A). Binding of PA28 to 20S opens the proteasome gate to allow entry of unfolded polypeptides, and because the sample is the endogenous complex pulled directly from living cells, it is likely that the density in the 20S antechamber represents unfolded polypeptides that have entered into the proteasome. Indeed, our cryo-EM map of double-capped PA28-20S-PA28 from the same sample shows additional density in both antechambers of 20S.

The observation of density in the antechamber of the PA28-20S complex suggests that the diffusion of unfolded polypeptides into the PA28-20S proteasome is a relatively fast process while the diffusion of polypeptides from the 20S antechamber into the catalytic chamber is relatively slow. Examination of the restriction between the antechamber and the catalytic chamber reveals the presence of numerous hydrophobic and aromatic amino acids, which may be serving to control the movement of charged polypeptides from the antechamber into the catalytic chamber (Figure 4B). Our results suggest a model for protein degradation by PA28-20S whereby unfolded proteins move quickly into the proteasome and held in the antechamber, followed by slower diffusion of the polypeptide into the catalytic chamber for cleavage and degradation (Figure 4C). This model suggests that the upregulation of PA28-20S in response to oxidative stress would help to quickly remove potentially toxic unfolded proteins from the cytoplasm followed by their subsequent degradation. Interestingly, no density is observed in the antechambers of 20S in any of our other proteasome cryo-EM maps, suggesting that this mechanism of peptide loading and degradation may be unique to the PA28-20S proteasome.

**Figure 4.**
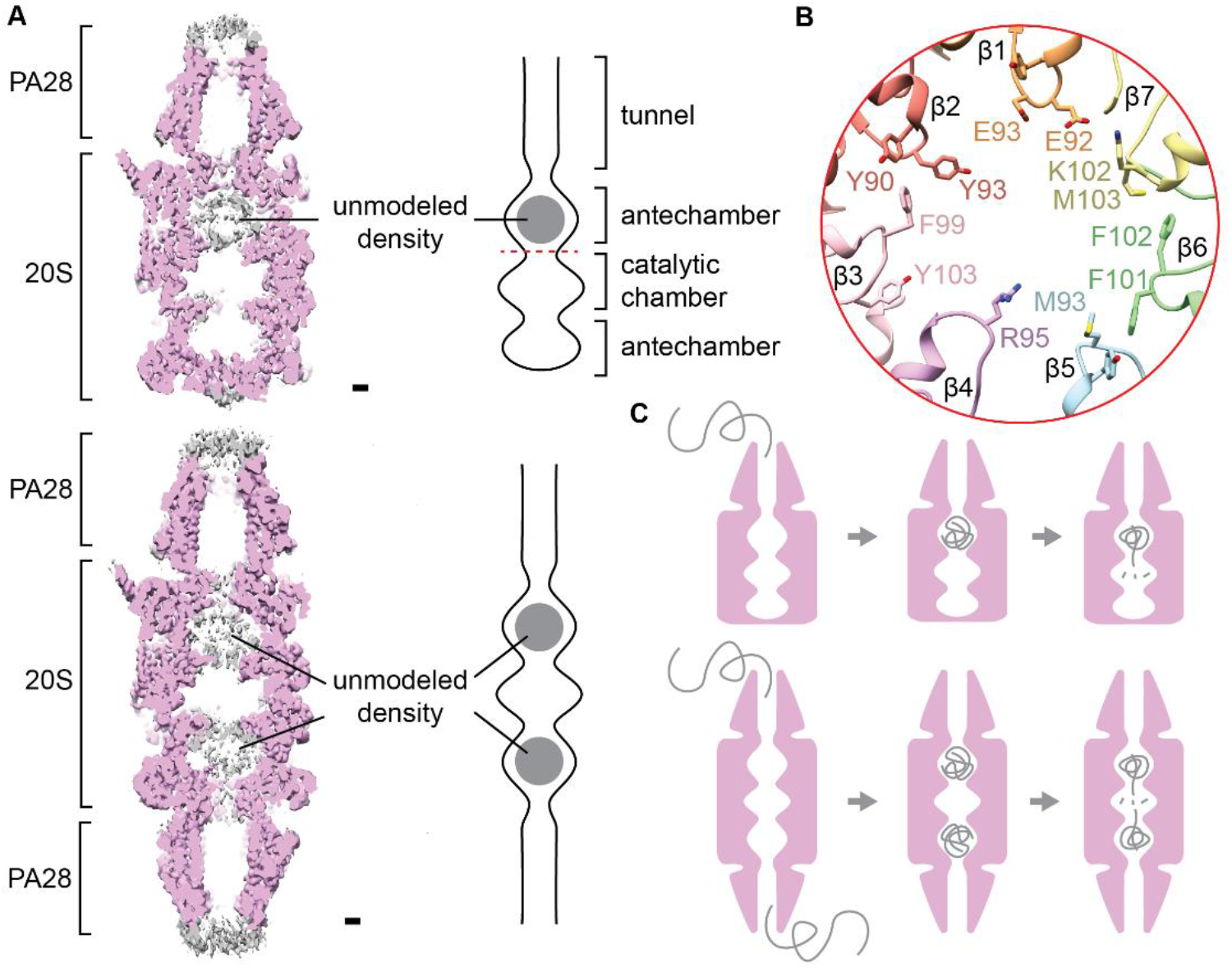
Polypeptide loading into PA28-20S and PA28-20S-PA28 proteasomal complexes. (**A**) Cross-section through the cryo-EM map of PA28-20S shows the presence of additional density inside the antechamber of 20S that is adjacent to PA28. The additional unmodeled density appears disordered and likely represents unfolded polypeptides loaded into the proteasome complex. While only one antechamber is filled in the single-capped PA28-20S assembly, both antechambers of 20S are filled in the double-capped PA28-20S-PA28. Scale bar, 10 Å. (**B**) The restriction between the antechamber and catalytic chamber of 20S is lined with numerous hydrophobic and aromatic amino acids, which may serve to control the movement of unfolded polypeptides from the antechamber into the catalytic chamber. (**C**) Model of polypeptide loading into the proteasome via PA28. Unfolded polypeptides transit quickly through PA28 and into the antechamber of 20S, where they are held prior to degradation. The polypeptides diffuse with slower kinetics into the catalytic chamber of 20S where they are subsequently degraded.

## DISCUSSION

Many proteins do not behave well when overexpressed in heterologous expression systems, which have made them refractory to structural studies. The approach demonstrated here provides a way to overcome these sample production limitations by enabling the rapid tagging of endogenous proteins in essentially any cell line. Not only does this approach simplify the study of important protein complexes, it also allows the purification of proteins in their cell-specific states. Combining this strategy with disease-relevant cell lines can allow the visualization of proteins in their disease states. These kinds of studies could help us better understand the mechanisms of protein regulation and how proteins become dysregulated in a wide range of maladies.

The targeting and purification of endogenous proteins can co-purify with their native interaction partners as demonstrated with the proteasome in this study. As a model system, prior knowledge about the complexes informed the tagging strategy, including targeting the C termini of the 20S β subunits and the N terminus of the PA28 α-subunit. For cases without prior structural knowledge, the highly efficient CRISPR/Cas knock-in pipeline in combination with FACS described here enables a shotgun approach of tagging multiple subunits in parallel. The approach described here is well suited for the study of the protein interactome, whereby a technique such as mass spectrometry can provide a big picture overview of a protein-protein interaction network while cryo-EM can zoom in and provide a high-resolution view of these interactions. Additionally, cryo-EM offers the ability to classify protein images and quantify the distribution of protein states, enabling insights into how proteins and protein complexes may change in response to different kinds of stress and cellular conditions. One current limitation in this approach is the potential for protein assemblies to dissociate during cryo-EM sample preparation, which would make it challenging to visualize weak protein-protein interactions. One possible solution would be to cross-link the protein complexes prior to imaging. In addition, improved computational methods to sort heterogenous protein images would also help to identify weak or rare protein-protein interactions in cryo-EM.

Most endogenous proteins exist at much lower concentrations in cells relative to overexpressed proteins, which can make both fluorescence selection and structural analysis of these relatively rare targets challenging. For selection, it has been estimated that ~ 50% of the human proteome is expressed above the FACS detection limit using mNG2[11] tagging ^14^, and signal amplification using our recent tag-assisted split enzyme complementation method ^26^ could further expand the accessible range. For purification, the recent development of affinity grids offers another potential solution to the study of low abundance targets ^27,28^. Tagging endogenous proteins and purifying them directly on an EM grid can drastically reduce the amount of sample required for structural studies by avoiding intermediate steps ^27^. Additionally, pulling down proteins directly from lysate can reduce the time the protein spends outside the cell environment, which can help to preserve their native protein-protein interactions and the integrity of delicate protein complexes.

One exciting future direction stemming from the concepts described here is the study of endogenous proteins from animal models. This can allow the structural analysis of proteins from their physiological contexts, which could open new avenues in the study of aging and neurodegenerative disorders. While the study of proteins from native sources offers many exciting prospects for structural biology and beyond, challenges remain. Improved methods to tag endogenous proteins with low expression and optimized strategies to target native proteins from animal models will be critical areas for future development.

## METHODS

### Primers for guide RNA (gRNA) synthesis

ML557: TAATACGACTCACTATAG

ML558: AAAAAAAGCACCGACTCGGTGC

ML611: AAAAAAAGCACCGACTCGGTGCCACTTTTTCAA GTTGATAACGGACTAGCCTTATTTAAACTTGCT ATGCTGTTTCCAGCATAGCTCTTAAAC

<u>PSMB2</u> oligo: TAATACGACTCACTATAG<u>GATGTTAGGAGCCCTGTTTG</u>GTTTAAGAGCTATGCTGGAA

<u>PSMB6</u> oligo: TAATACGACTCACTATAG<u>TTATTGCATACTAGAATCCC</u>GTTTAAGAGCTATGCTGGAA

<u>PSMB7</u> oligo: TAATACGACTCACTATAG<u>GCCACCCACTGATGC CATTC</u>GTTTAAGAGCTATGCTGGAA

<u>PSME1</u> oligo: TAATACGACTCACTATAG<u>CCCGGTCATGGCCAT GCTCA</u>GTTTAAGAGCTATGCTGGAA

### Single strand DNA for homology-directed repair (HDR)

PSMB2-<u>mNG211</u>-<u>StrepII</u>: ATTGACAAAAATGGCATCCATGACCTGGATAAC ATTTCCTTCCCCAAACAGGGCTCCGGCAGC<u>ACCGAGCTCAACTTCAAGGAGTGGCAAAAGGCCT TTACCGATATGATG</u>GGGAGTGCG<u>TGGTCCCACCCTCAATTCGAGAAG</u>TAACATCATGTCCTCCCTCCCACTTGCCAGGGAACTTTTTTTTGATGGGCTCCTTT

PSMB6-<u>mNG211</u>-<u>StrepII</u>: CAAGTACTTTTGGGAGACCAGATACCCAAATTCGCCGTTGCCACTTTACCACCCGCCGGCAGC<u>ACCGAGCTCAACTTCAAGGAGTGGCAAAAGGCCTTTACCGATATGATG</u>GGGAGTGCG<u>TGGTCCCACCCTCAATTCGAGAAGTG</u>AATACTGGGATTCTAGTATGCAATAAGAGATGCCCTGTACTGATGCAAAATTTA

PSMB7-<u>mNG211</u>-<u>StrepII</u>: ATCACTCCTCTGGAGATTGAGGTGCTGGAAGAAACAGTCCAAACAATGGACACTTCCGGCAGC<u>ACCGAGCTCAACTTCAAGGAGTGGCAAAAGGCC TTTACCGATATGATG</u>GGGAGTGCG<u>TGGTCCCACCCTCAATTCGAGAAG</u>TGAATGGCATCAGTGG GTGGCTGGCCGCGGTTCTGGAAGGTGGTGAGCATTGAGGC

<u>ALFA</u>-<u>sfCherry211</u>-PSME1: CCCACTCCACTCCTTGTGCGGCGCTAGGCCCC CCGTCCCGGTCATG<u>CCTTCTAGATTAGAAGAGGAATTGAGACGACGATTGACTGAACCA</u>GGATC<u>TTATACTATCGTTGAACAATACGAGAGAGCTGA GGCAAGACACTCTACA</u>GGTTCTGGCGCCATGC TCAGGGTCCAGCCCGAGGCCCAAGCCAAGGTGAGCGCCG

### gRNA IVT Template Synthesis

The IVT template for LMNA gRNA was made by PCR (Supplementary Figure 1). The reactions were completed in a 100μL reaction containing 50μL 2x Phusion MM, 2 μL ML557+558 mix at 50 μM, 0.5 μL ML611 at 4 μM, 0.5 μL of each gene-specific oligo at 4 μM, and 47 μL DEPC H20. PCR program: 95°C, 3 min; 20 X (98°C, 20 s; 57°C, 15 s; 72°C, 5 s); 72°C, 60 s. The PCR product was purified using a Zymo DNA Clean and Concentrator Kit (Zymo Research D4013).

### gRNA Synthesis

IVT was carried out using the HiScribe T7 Quick High Yield RNA Synthesis Kit (NEB E2050S) with the addition of RNAsin (Promega N2111). Purification of mRNA was performed using the RNA Clean and Concentrator Kit (Zymo Research R1017). gRNA was stored at −80C immediately after measuring concentration and diluting to 130 μM.

### Cas9 HDR Knock-Ins

Ribonucleoproteins (RNP’s) were generated in 40 μL reactions consisting of 4μL of sgRNA at 130 μM, 10μL of purified Cas9 at 40 μM, 6μL of HDR template at 100 μM, 8μL of 5× Cas9 Buffer, and DEPC H2O up to 40 μL. HDR template ultramer sequences synthesized from IDT are provided above.

In a sterile PCR or microcentrifuge tube, Cas9 Buffer, DEPC H2O, and sgRNA were mixed and incubated at 70 °C for 5 min to refold the gRNA. During this step, 10 μL aliquots of purified Cas9 at 40 μM were thawed on ice and once thawed were slowly added to the diluted sgRNA in Cas9 buffer and incubated at 37 °C for 10 min for complex formation.

Finally, 6μL of each ultramer donor was added to the RNP mix, and all samples were kept on ice until ready for nucleofection. For efficient recovery post-KI, a 6-well plate with 2mL of Expi293F expression media (ThermoFisher) per well was incubated at 37°C. An appropriate amount of supplemented Amaxa solution corresponding to the number of KIs to be performed was prepared at room temp in the cell culture hood. For each sample, 65.6 μL of SF solution and 14.4 μL of supplement was added to an Eppendorf tube for a total of 80 μL per KI. Amaxa nucleofector instruments/computers were then turned on and kept ready for nucleofection.

Expi293F cells were harvested into a sterile tube and counted. A volume equivalent to 800k cells per KI was transferred to another tube and centrifuged at 500g for 3 min. Supernatant was removed and cells were resuspended in 1 mL of PBS to wash. The cells were centrifuged again at 500g for 3 min. PCR tubes containing RNPs were brought into the TC hood. Cells were resuspended in supplemented Amaxa solution at a density of 10k cells/μL. 80μL of the cell resuspension was added to each 10μL RNP tube. The cell/RNP mix was pipetted into the bottom of the nucleofection plate. The nucleofection was carried out in cuvettes on a Lonza Nucleofector X Unit (Lonza AAF-1002X) attached to Lonza 4D Nucleofector Core Unit (Lonza AAF-1002B). Cells were nucleofected using FS-100 program and recovered using 400 μL of media from the prewarmed 6-well plate and transferred to the corresponding well. The cells were incubated at 37 °C and 8 % CO_2_.

Once cells expanded to the point where they began to detach from the well bottom, the plate was transferred to a shaker operating at 120 RPM and incubated at 37 °C and 8 % CO_2_. The next day, the cells were transferred to a 125 ml vented flat-bottom Erlenmeyer flask containing 20 ml Expi293F media and incubated at 37 °C and 8 % CO_2_ with shaking at 120 RPM until the cell density reached 1 million/ml.

### Transfection of mNG2(1-10)

To prepare transfection mixture, 20 ug of DNA plasmid containing mNG21-10 was added to 1 ml of Opti-MEM (Gibco) in one tube and 53 ul of ExpiFectamine 293 (Gibco) was added to 1 ml of Opti-MEM (Gibco) in a separate tube. The mixture was incubated at room temperature for 5 mins. The Opti-MEM containing Expifectamine 293 was then added to the solution containing DNA, inverted multiple times to mix, and then incubated at room temperature for 25 mins. The transfection mixture was then added to 20 million Expi293F cells in 20 ml of media in a 125 ml vented flat-bottom Erlenmeyer flask. The cells were incubated at 37 °C for 2 days with shaking at 120 RPM.

### Flow Cytometry Analysis and FACS

FACS sorting and flow cytometry was performed on a BD FACSAria II in the Laboratory for Cell Analysis at UCSF. mNG2 signal was measured with the 488 nm laser and 530/30 bandpass filter. 1 million cells/sample were sorted into Eppendorf tubes and then plated in a 6-well plate. The 6-well plate was incubated at 37 °C and 8 % CO_2_ for 2 days with shaking at 120 RPM.

### cDNA Analysis

Total RNA was extracted from 1 million cells using the Monarch Total RNA Miniprep Kit (NEB, #T2010S). We prepared cDNA from 1μg of extracted RNA using LunaScript^®^ RT SuperMix Kit (NEB, #E3010). No Template and No Reverse Transcriptase controls (NTC and NRT) were performed in parallel to cDNA preparations. Sequencing confirmation of amplicons was completed by Elimbio.

### Cell culture and protein purification

Expi293F cell lines were grown in 100 ml Expi293 expression medium (ThermoFisher) in 500 ml flat bottom Erlenmeyer flasks at 37°C, 8% CO_2_, and shaking at 120 RPM. When cells reached a density of 3 million cells per ml, cells were harvested by centrifugation at 1000g for 5 mins. Cells were resuspended in 20 ml Buffer A (50 mM HEPES, 150 mM NaCl, 1 mM DTT, pH 7.5) supplemented with SigmaFast protease inhibitor (Sigma). Cells were lysed by sonication and insoluble debris were removed by ultracentrifugation at 100,000g for 20 mins. For StrepII-tagged proteins, the supernatant was passed through a column containing 0.2 ml Strep-Tactin XT Sepharose resin (Cytiva) by gravity flow. The resin was washed with 2 ml Buffer A and the bound proteins were eluted with 1 ml Buffer A supplemented with 50 mM Biotin (final concentration). For ALFA-tagged proteins, the supernatant was passed through a column containing 0.2 ml ALFA Selector PE resin (NanoTag Biotechnologies) by gravity flow. The resin was washed with 2 ml Buffer A and the bound proteins were eluted at room temperature with 1 ml Buffer A supplemented with 200 μM ALFA peptide (final concentration). To acquire 19S-bound proteasome complexes, 1 mM AMPPNP (Sigma) and 2 mM MgCl was included throughout the purification. The eluted proteins were concentrated in a 100K centrifugal concentrating device.

### Negative Stain TEM Imaging

Negative stain EM grids were prepared following established protocol ^29^. Specifically, 3.5 μL of purified proteasome complexes at 0.02 mg/ml was pipetted onto a continuous carbon coated grid and incubated at room temperature for 60 s. The sample was blotted using filter paper. Immediately, 3.5 μL of 2% (w/v) uranyl formate was pipetted onto the grid and blotted away with filter paper. This treatment with uranyl formate stain was repeated two more times, except an addition of a 30 s wait between stain application and blotting during the last cycle. The grid was left to dry at room temperature for 2 mins. The grids were imaged in a FEI Tecnai T12 fitted with a Gatan US4000 CCD camera. The images were processed using CryoSPARC ^30^.

### Cryo-EM Imaging and data processing

To prepare sample grids, 3.5 μL of purified proteasome complexes at 2 mg/ml was applied to a Quantifoil Au 300 mesh 1.2/1.3 grid in a Vitrobot Mark IV (Thermo), blotted, and plunge frozen in liquid ethane. The sample grids were imaged in a FEI Polara equipped with field emission source and operated at 300kV. Cryo-EM data acquisition is performed by using SerialEM ^31^. Images were recorded using a K2 Summit electron detector operated in super-resolution mode using 8 electrons/pixel/second at 1.22 Angstroms/physical pixel. The images were corrected for specimen drift using MotionCor2 ^32^. Non-dose weighted sums were used for CTF determination using gCTF, and dose weighted sums were used for the rest of single particle cryo-EM image processing. Particle picking, classification and refinement were processed in CryoSPARC ^30^ and cisTEM ^33^. Atomic models were built into the cryo-EM density maps using Coot ^34,35^ and Phenix ^36,37^. The resulting cryo-EM maps and models were visualized using UCSF Chimera ^38^.

## Data Availability

Cryo-EM density maps and atomic models reported in this study have been deposited in the Electron Microscopy Data Bank (EMDB) and Protein Data Bank (PDB) under the accession codes EMD-24275 and PDB 7NAN (20S), EMD-24276 and PDB 7NAO (PA28-20S), EMD-24277 and PDB 7NAP (PA28-20S-PA28), and EMD-24278 and PDB 7NAQ (PA200-20S).

## Acknowledgement

This work is supported by grants from National Institute of Health (R35GM140847 and P50AI150476 to Y.C. and R01GM124334 and R01GM131641 to B.H). Instruments at UCSF Cryo-EM facilty are partially supported by grants from National Institute of Health (S10OD020054, S10OD021741 and S10OD026881) and Howard Hughes Medical Institute. B.H. is a Chan Zuckerberg Biohub investigator. Y.C. is an Investigator of Howard Hughes Medical Institute.

## Author contribution

S.M. designed the Cas9 guide RNA and DNA, and performed the CRISPR knock-in and FACS. J.Z. prepared the cells and carried out protein purification, cryo-EM data acquisition and processing. J.Z., S.M., B.H and Y.C. conceived the project, interpreted the results and wrote the manuscript.

