## supplementary_figures for "Efficient tagging and purification of endogenous proteins for structural studies by single particle cryo-EM"

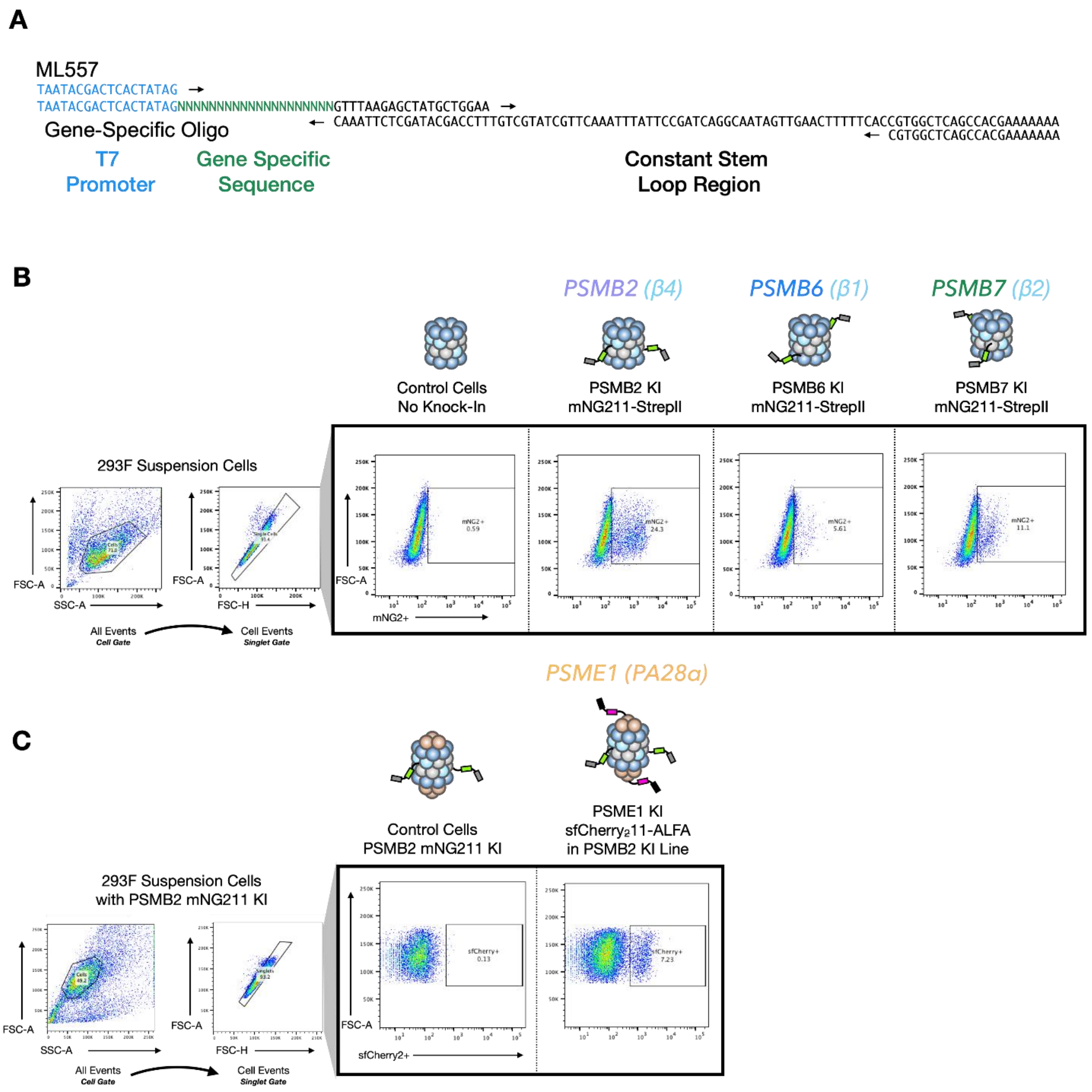


**Supplementary Figure 1. gRNA IVT Template Design and FACS Sorting of KIs.** (**A**) PCR schematic for the IVT template to make Cas9 gRNA. The gene specific primers (provided in the Methods section) anneal to ML611 containing the constant stem loop region. ML557 and ML558 are used to amplify the gene-specific template created by the gene-specific oligo and ML611. (**B**) FACS sorting of PSMB knock-in Expi293F cells. Shown are control cells used to set gating and PSMB2, PSMB6, and PSMB7 KIs containing the mNG211-StrepII donor. Cells in the gate were collected for downstream structural studies. (**C**) FACS sorting of PSME1 knock-in. This knock-in was performed in the sorted PSMB2 KI cell line. Shown are control cells used to set gating and PSME1 KI containing the sfCherry$_2$11-StrepII donor. Cells in the gate were collected for downstream structural studies.

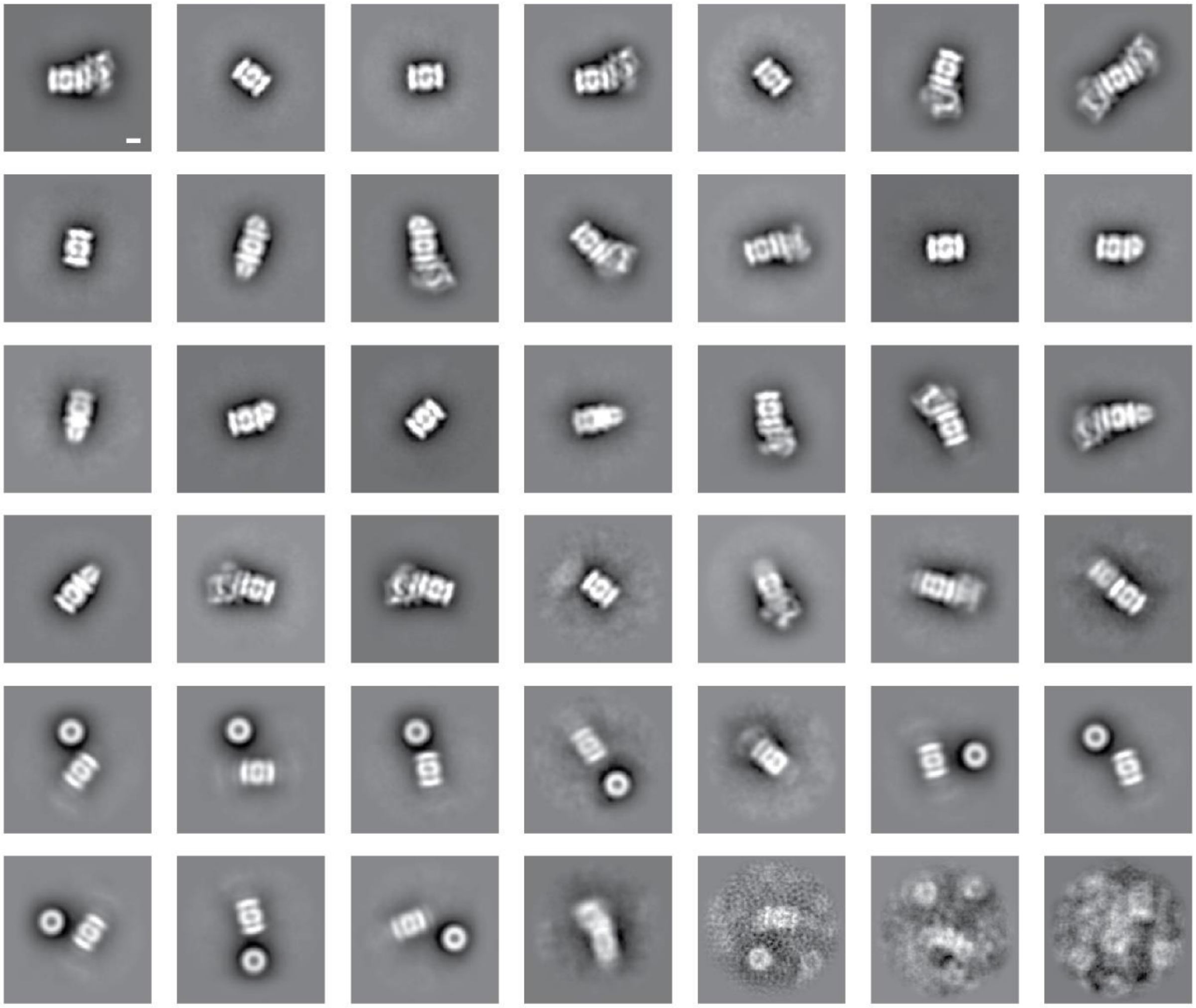

**Supplementary Figure 2. Negative stain EM of β4-tagged proteasomal complexes.** 2D classification of the images show a diverse population of different proteasomal complexes formed between 20S, 19S, PA28, and PA200. Scale bar, 50 Å.

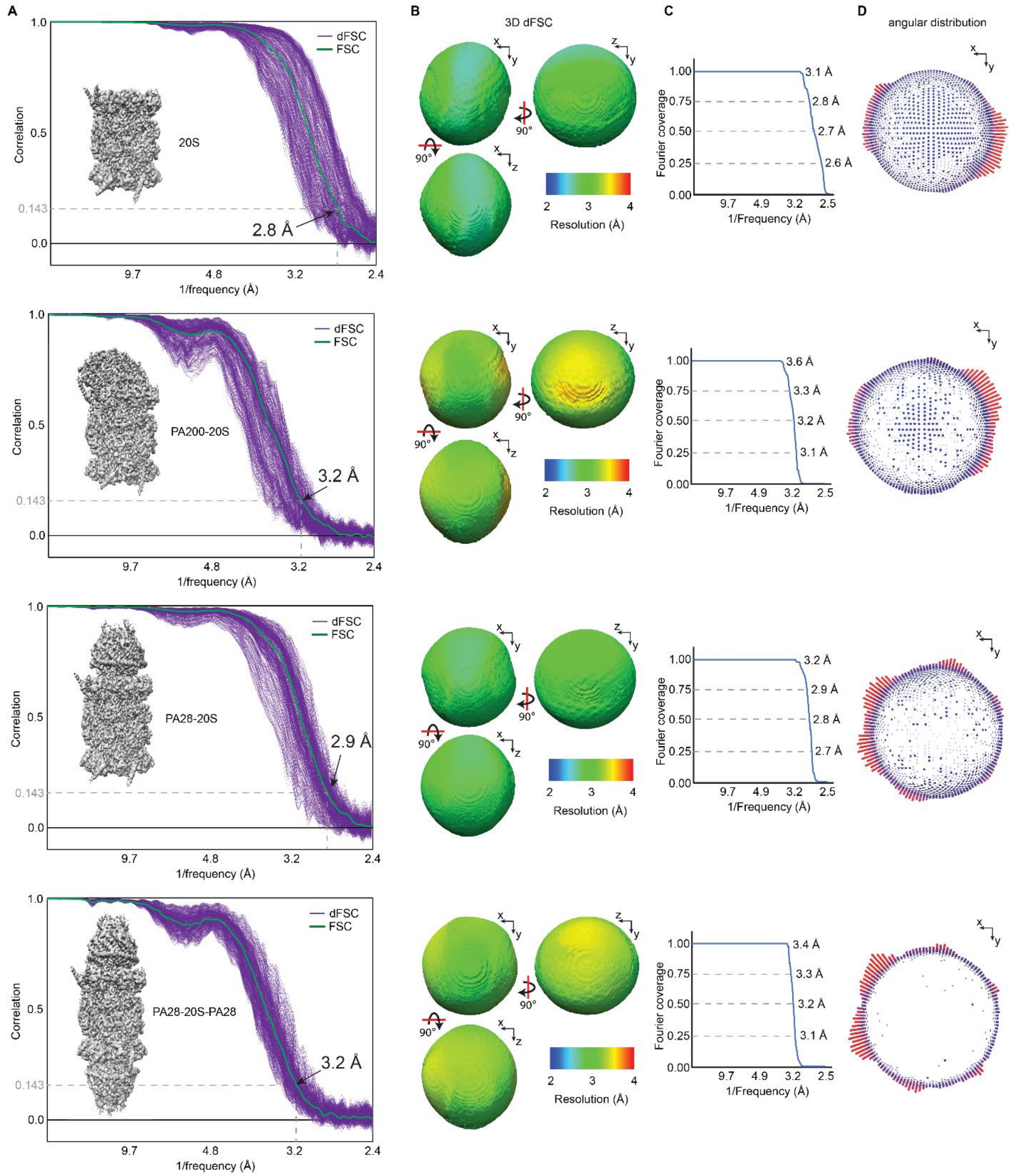


**Supplementary Figure 3. Cryo-EM map statistics.** (**A**) Fourier shell correlation (FSC) and 1D directional FSC (dFSC) plots. (**B**) Visualization of 3D dFSC. (**C**) Plots showing coverage of Fourier space. (**D**) Plots showing distribution of viewing angles from final refined datasets.

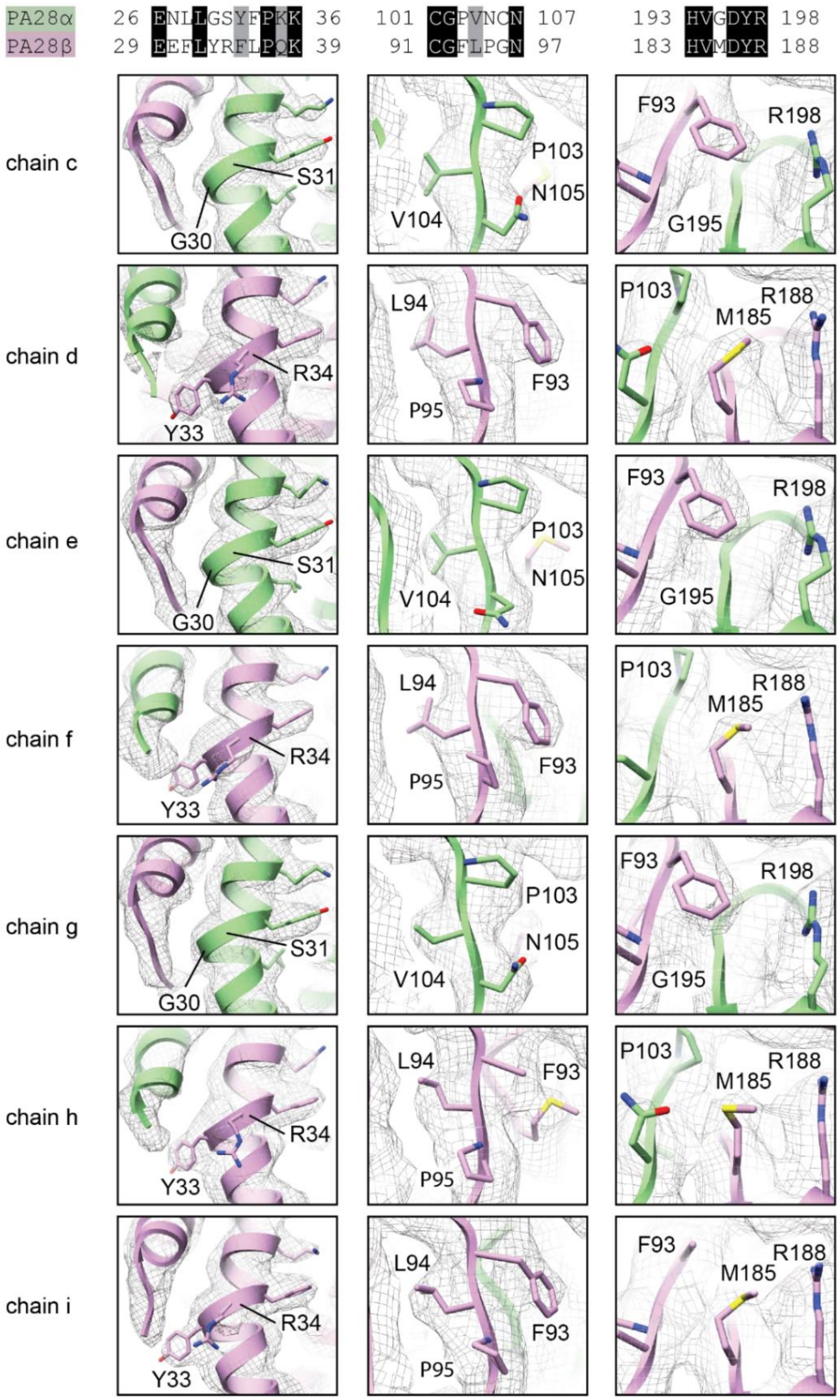


**Supplementary Figure 4. Fit of PA28 model in cryo-EM density**. Residue pairs that distinguish between PA28 α/β subunits include G30/Y33, S31/R34, P103/F93, and G195/M185.

**Supplementary Table 1. Cryo-EM imaging parameters**

| | | | | |
|---|---|---|---|---|
| Microscope | Polara (300 kV) | | | |
| Camera | Gatan K2 Summit | | | |
| Exposure rate | 8 e/pixel/s | | | |
| Total exposure | 43 e/Å^2 | | | |
| Pixel size | 1.22 Å/pixel | | | |
| Micrographs | 3761 | | 5994 | |
| Class | 20S | PA200-20S | PA28-20S | PA28-20S-PA28 |
| Particle images | 499,629 | 50,767 | 135,937 | 22,946 |
